# Identifying Fundamental Gaps in Functional Metagenomics: A Step Towards Unlocking Microbiome Research Potential

**DOI:** 10.64898/2026.01.10.698778

**Authors:** Sumeet Kumar Tiwari, Andrea Telatin, Dipali Singh

## Abstract

Incomplete functional annotation systematically limits biological interpretation in microbiome studies and their translational potential. Poor annotation arises from multiple causes1 with incomplete gene-protein-reaction mapping being one tractable yet under-examined contributor. We address this gap by developing a comprehensive hierarchical framework that systematically integrates gene families in UniRef1 proteins in UniProt1 and metabolic reactions in BioCyc1 mapping 76% of reactions (432,510 of 562,1869 reactions in BioCyc across 211244 organisms) to the encoding gene families using EC and Pfam domain-based strategies. To demonstrate how database mapping choices impact metagenomics studies1 we applied our framework to human gut metagenome dataset using HMP Unified Metabolic Analysis Network (HUMAnN) and compared outcomes with HUMAnN’s default database mapping approach. Our GPR mapping increased detectable metabolic reactions by 28-fold and significantly increased the data prevalence across samples (from 35% to 80% core reactions). This directly addresses the problem of data sparsity1 a critical barrier to statistical and machine learning applications. These gains derive from systematic database integration alone1 without predictive algorithms1 demonstrating that substantial functional dark matter and data sparsity problems in microbiome studies arises from methodological artifacts that are directly addressable.

## 1 Introduction

The study of microbial communities has traditionally relied on taxonomic profiling to determine *who is present*, using phylogenetic markers or genome sequencing [1]. However, understanding microbial contributions to host physiology, disease, or ecosystem processes requires insight into *what they do*: the gene products, biochemical activities, and metabolic capabilities that mediate interactions with their environment or host [2]. This functional perspective is foundational to translating microbiome research into clinical diagnostics, therapeutic interventions, and biotechnological applications [3, 4]. Modern metagenomic studies increasingly recognise that taxonomic and functional perspectives are complementary. While taxonomy defines community structure and biogeographical patterns [5], functional profiling reveals metabolic potential, biochemical activity, and ecosystem-level functions [6, 7]. Integrating these perspectives links taxa to specific functions, identifies functional redundancy, and provides a mechanistic basis for assessing community metabolic capacity which is a critical requirement for personalised medicine approaches that leverage microbiome modulation [8, 9].

Microbiome studies have widely converged on metabolic pathways as the primary metric of functional profiling [6, 10]. While pathways offer dimensionally reduced representations of metabolic function, they are modular constructs that do not fully capture cellular function. In MetaCyc (version 25.1), a well-curated metabolic pathway database, only 63.8% of reactions carry pathway annotations, meaning one-third of known enzymatic functions are systematically excluded during pathway-based functional profiling. Therefore, a comprehensive functional profiling requires moving beyond pathway-centric approaches toward multi-resolution functional profiling.

At the core of such multi-resolution profiling lies the gene-protein-reaction (GPR) relationship. Despite the importance of comprehensive GPR mapping for functional characterisation, substantial fractions of metagenomic data remain functionally uncharacterised. This “functional dark matter” arises from multiple causes, including genuinely unknown gene functions but also incomplete gene-protein-reaction mapping and cross-database integration that are tractable yet systematically overlooked contributors. Understanding the extent and consequences of this database integration gap is essential for improving functional annotation coverage and enabling more comprehensive metabolic characterisation of microbial communities.

To address this gap, we developed a comprehensive, hierarchical GPR mapping framework that integrates BioCyc’s extensive reaction database (562,869 reactions) [11] with UniRef90 gene families [12] through both EC and protein domain (Pfam [13])-based mapping strategy. This modular approach enables functional annotation at multiple resolutions, from individual genes to proteins and enzymatic reactions, allowing researchers to select the appropriate functional granularity for their specific research questions. The framework can be applied directly as custom mapping files with existing functional metagenomics tools such as HMP Unified Metabolic Analysis Network (HUMAnN) [6], or integrated with outputs from genome annotation pipelines such as BAKTA where UniRef90 gene identifiers are available. We applied our mapping framework to human gut metagenome dataset using HUMAnN and compared outcomes with HUMAnN’s default database mapping approach at reaction resolution. Our GPR mapping approach significantly increased detectable metabolic reactions, and their prevalence across samples providing basis for more robust comparative analyses in microbiome studies.

## 2 Methods

### 2.1 Database Sources and Versions

We integrated data from three major biological databases to construct comprehensive gene-protein-reaction mappings as shown in Figure 1. UniProtKB [14] (version 2019_01, including Swiss-Prot and TrEMBL) and UniRef90 [12] (version 2019_01, clustered at 90% sequence identity) were obtained from the UniProt Consortium. BioCyc [11] (version 28.5, December 10, 2024), a collection of organism-specific pathway / genome databases covering thousands of organisms across all domains of life, was obtained under the appropriate license from SRI International.

**Figure 1:**
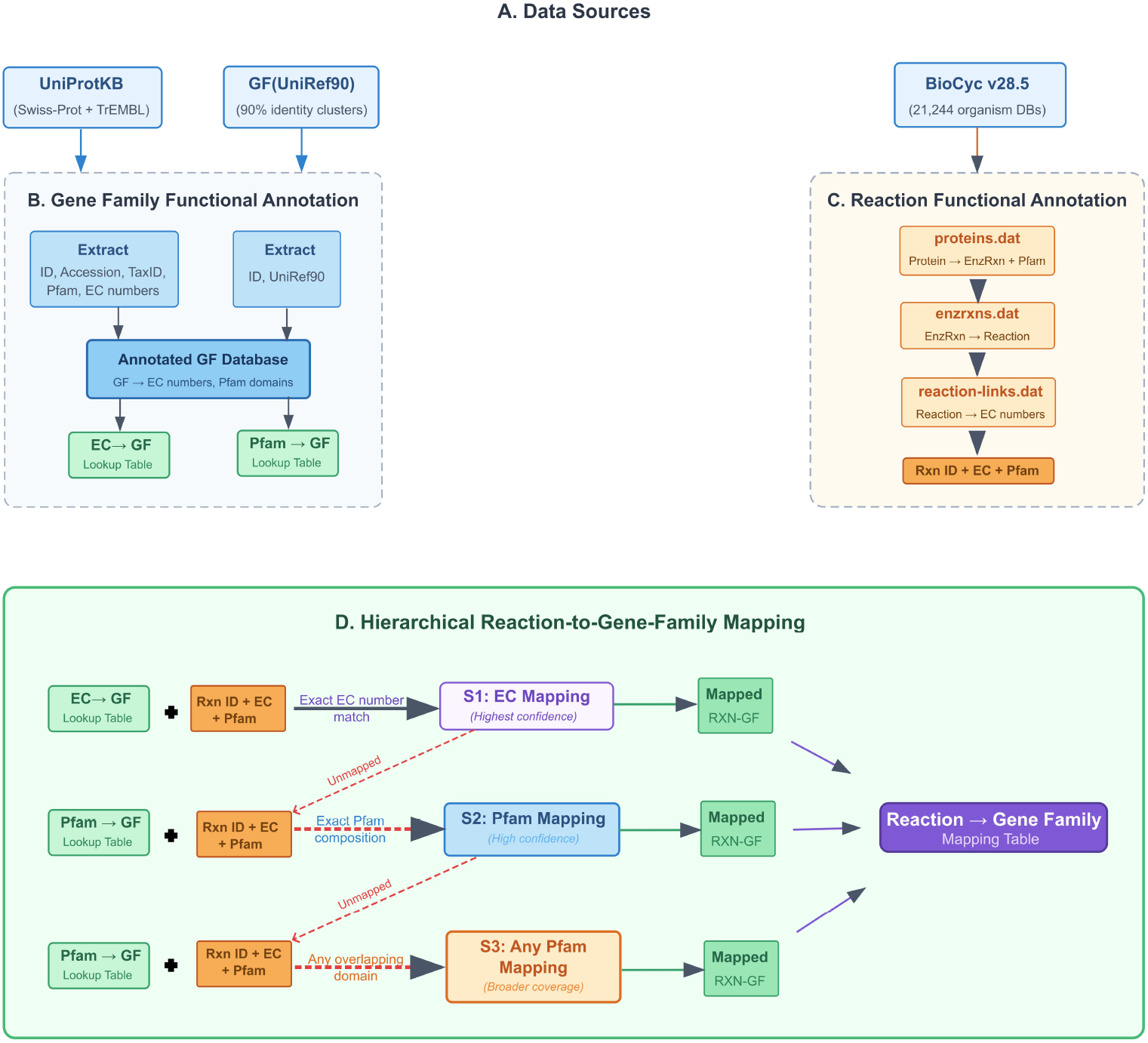
Database integration workflow. Systematic integration strategy connecting UniRef90 gene families, UniProt protein annotations, and BioCyc metabolic reactions through EC-based and Pfam domain-based hierarchical mappings.

### 2.2 Gene Family to Protein Mapping

Protein-level annotations were obtained from UniProtKB by parsing both Swiss-Prot and TrEMBL databases. A unified annotation table containing unique identifier (ID), protein accessions, NCBI taxonomy identifier, Pfam domain(s), and EC number(s) was created. These annotations were subsequently linked to UniRef90 gene families by extracting the clusters’ membership information from the UniRef90 (.xml) database. These gene families were then annotated with EC number(s) and Pfam domain(s) inherited from their constituent protein members, creating a comprehensive gene-family (GF) annotated database.

To facilitate downstream reaction mapping, we constructed two complementary lookup tables: (1) an EC-to-GF mapping by grouping all UniRef90 clusters sharing the same EC number, and (2) a Pfam-to-GF mapping by grouping UniRef90 clusters with identical Pfam domain composition.

### 2.3 Protein to Reaction Mapping

BioCyc version 28.5 comprises 21,244 organism-specific databases, which were used to obtain reaction-level functional information. For each organism-specific database, a multi-step parsing workflow was implemented. First, protein-to-enzymatic-reaction relationships and their associated Pfam domain(s) were extracted from `proteins.dat.` Second, enzyme-to-reaction mappings were retrieved from `enzrxns.dat.` Finally, EC number(s) annotations were incorporated from `reaction-links.dat`, resulting in organism-specific reaction annotation files containing reaction identifiers together with their associated EC numbers and Pfam domains.

### 2.4 Hierarchical Reaction-to-Gene Family Mapping Strategy

We employed a hierarchical three-tier mapping strategy to link BioCyc reactions to UniRef90 gene families, proceeding from high-confidence to broader coverage in three sequential steps. Each step processes only reactions that remained unmapped in previous steps, ensuring non-redundant assignments.

#### Step 1: EC-based mapping

Reactions with EC number annotations were mapped to gene families sharing the same EC number using the EC-to-gene-family lookup table. This provides the highest-confidence mappings as EC numbers directly encode enzymatic function.

#### Step 2: Pfam-based mapping

For reactions that could not be assigned to gene families in Step 1, Pfam domain information was used as an alternative mapping strategy. Reactions were linked to gene families sharing identical Pfam domain compositions, capturing proteins that are well-curated at the domain level but lack formal EC classification.

#### Step 3: Relaxed Pfam-based mapping

For remaining unmapped reactions with Pfam annotations, we relaxed the matching criterion to allow mapping when *any* Pfam domain from the reaction annotation matched *any* domain in a gene family’s annotation. This maximises coverage but could introduce lower specificity.

### 2.5 Functional Profiling of Shotgun Metagenome

Functional profiling of shotgun reads was performed by HUMAnN3 (v3.6.0) [15]. HUMAnN3 internally executes MetaPhlAn (v3.1.0) to infer species-level taxonomic composition based on clade-specific marker genes, using the `mpa _v30_CHOC0PhlAn_201901` marker database [15]. Based on the detected taxa, HUMAnN3 constructed a sample-specific pangenome database from ChocoPhlAn and aligned reads to these species-specific reference sequences to identify gene families at the nucleotide level. Reads that did not map to the species-specific pangenomes were subsequently translated and searched against the UniRef90 protein database (`uniref90_201901b`) using DIAMOND to recover additional gene-family assignments. Gene-family abundances were reported by HUMAnN3 in reads per kilobase (RPK), which normalizes gene length.

For each sample, UniRef90 gene-family abundance profiles were converted to reaction-level profiles using the `humann_regroup_table` utility with two different mapping approaches: the default HUMAnN3 and our comprehensive mapping (section 2.4). At reaction resolution, reads were classified as UNMAPPED (not aligned to gene families), UNGROUPED (aligned to gene families but not assigned to metabolic reactions), or GROUPED (aligned to gene families and assigned to metabolic reactions).

Read counts were normalised to percentages of total reads for unmapped and percentages of mapped reads (total reads minus unmapped) for ungrouped and grouped categories. Distributions were visualised using boxplots with swarm plots showing individual samples. Statistical di erences between mapping approaches for ungrouped and grouped categories were assessed using paired Wilcoxon signed-rank tests, with significance thresholds at p < 0.001.

## 3 Results and Discussion

### 3.1 Comprehensive mapping of reactions to gene families

We used a hierarchical three-tier mapping strategy, as described in section 2.4 and Figure 1, to systematically map reactions in BioCyc, including MetaCyc, to their corresponding EC numbers and protein domains (Pfam), and subsequently linked these to UniRef90 gene families. Figure 2 summarizes the distribution of reactions captured at each mapping level across both BioCyc and MetaCyc databases. The latter is a reference metabolic pathway database within the BioCyc collection. MetaCyc-specific reaction—gene family associations were derived by restricting the BioCyc mappings to reactions present in MetaCyc.

**Figure 2:**
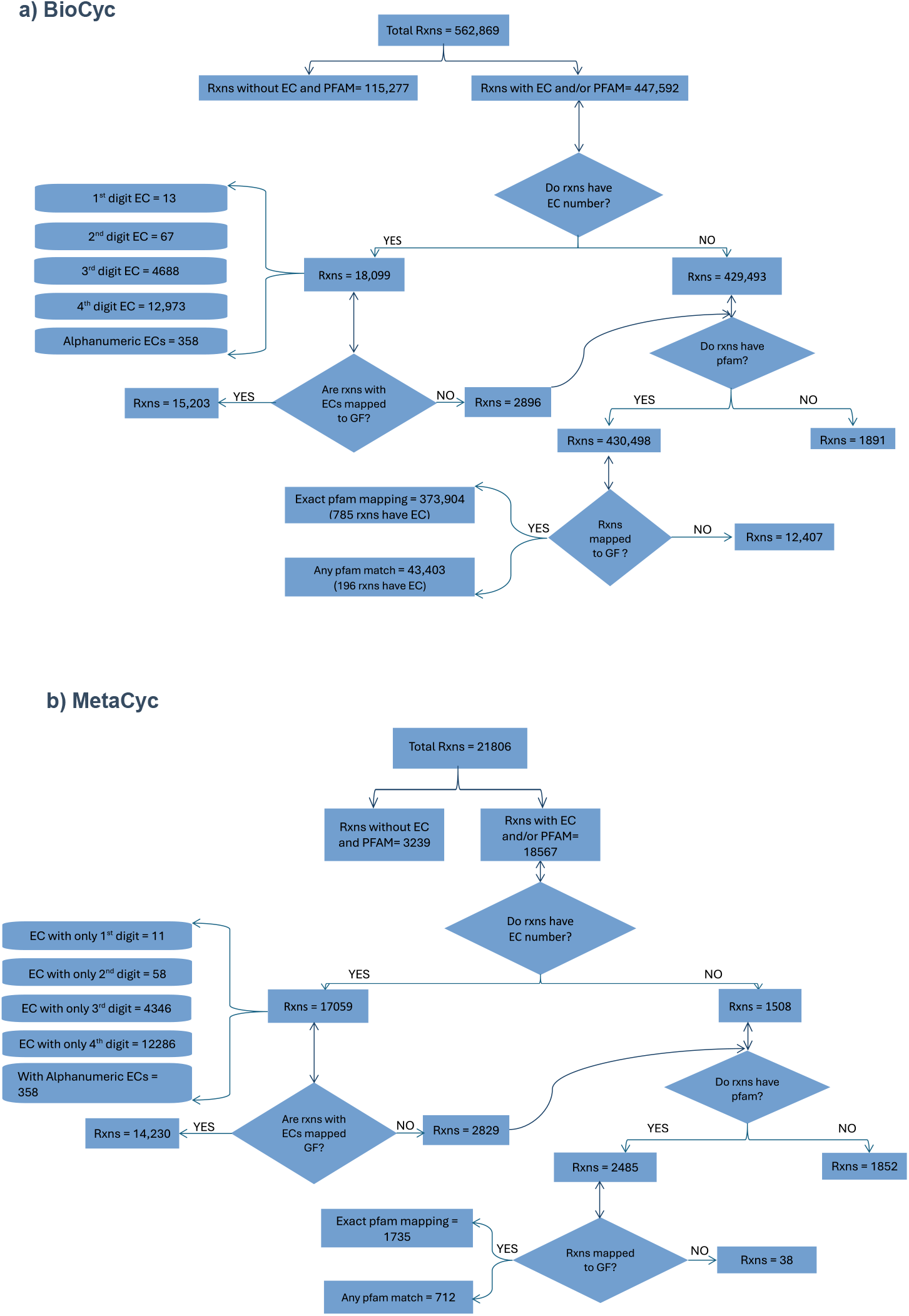
Database annotation coverage at each hierarchical mapping level. (A) Bio-Cyc database (562,869 reactions): Distribution by annotation type showing limited EC coverage (2.7%) versus extensive Pfam coverage (76.3%). (B) MetaCyc database (21,806 reactions): Even in this curated database, 43.7% of reactions lack EC annotations despite having protein associations.

#### BioCyc reaction coverage

As shown in Figure 2a, of the 562,869 reactions in BioCyc (version 28.5), approximately 20.5% (115,277 reactions) lacked both Pfam and EC annotations, precluding their assignment to gene families. Among these unmappable reactions, only 707 (0.61%) were annotated as spontaneous reactions not requiring enzymatic catalysis, suggesting that the vast majority represent annotation gaps rather than non-enzymatic chemistry. For the remaining 79.5% (447,592 reactions) with either Pfam and/or EC annotation, the coverage by EC numbers was remarkably limited: only 3.2% (18,099 reactions) had EC number, and a mere 2.3% (12,973 reactions) possessed complete four-digit EC classifications. In contrast, 76.3% of BioCyc reactions (429,493) are linked to protein domains but lack EC assignments, representing a substantial pool of biochemically characterised reactions that remain inaccessible to EC-only mapping approaches.

#### MetaCyc reaction coverage

As shown in Figure 2b, MetaCyc exhibits better EC coverage than BioCyc but still shows substantial annotation gaps. Of 21,806 MetaCyc reactions, 12,286 (56.3%) carry complete four-digit EC annotations, while 43.7% lack EC assignments despite many having protein-level associations. This highlights that even within curated, manually reviewed databases, EC-based mapping captures only a fraction of characterised metabolic reactions.

This gap is particularly relevant for functional metagenomics tools such as HUMAnN, which map MetaCyc reactions to gene families using a two-tier strategy: direct protein-based associations via UniProt accessions, complemented by EC-based transitive mappings based on complete four-digit EC numbers [6].

### Hierarchical gene family mapping results

From a reaction-centric perspective, as shown in Figure 2.2 7% of BioCyc reactions (15,203) mapped to gene families via EC numbers alone, while 74.1% (417,307 reactions) mapped via Pfam domains (including 66.4% through exact Pfam matches and the remainder through relaxed domain matching) n total, we established comprehensive Reaction-Protein-Gene associations for 76% of all BioCyc reactions (432,510 reactions), a 28-fold improvement over EC-only approaches.

From a gene-centric perspective, 47.8% of the 87,296,736 UniRef90 families (41,691,001) lacked both EC and Pfam annotations, likely representing hypothetical proteins or domains not yet characterised. Among annotated gene families, 7.61% carried both EC and Pfam assignments, 0.42% had EC only, and 44.2% had Pfam only. Through our hierarchical framework, 36. 2% of all UniRef90 families could be mapped to biochemical reactions: 4.15% via EC alone, 7.54% via both EC and Pfam, and 28.2% via Pfam alone. This demonstrates that Pfam-based mapping captures the majority of functionally annotatable gene families, far exceeding the coverage achievable through EC-based approaches alone.

### 3.2 Functional Profiling of Gut Microbiome: A Case Study

We applied our GPR mapping framework to human gut metagenomic datasets (BioProject PR-JNA749645) to assess how database integration approach affects functional profiles. Functional profiles at reaction resolution were generated using HUMAnN3 pipeline, as described in section 2.5.

As shown in Figure 3, the default HUMAnN3 output maps ≈ 35% of reads to function at reaction resolution. Our customised GPR mapping approach maps ≈ 45% of reads to function at reaction resolution. The percentage of reads assigned to genes with defined metabolic functions is significantly higher using our approach, reducing the fraction of reads classified as “ungrouped” (mapped to genes but without metabolic function assignments).

**Figure 3:**
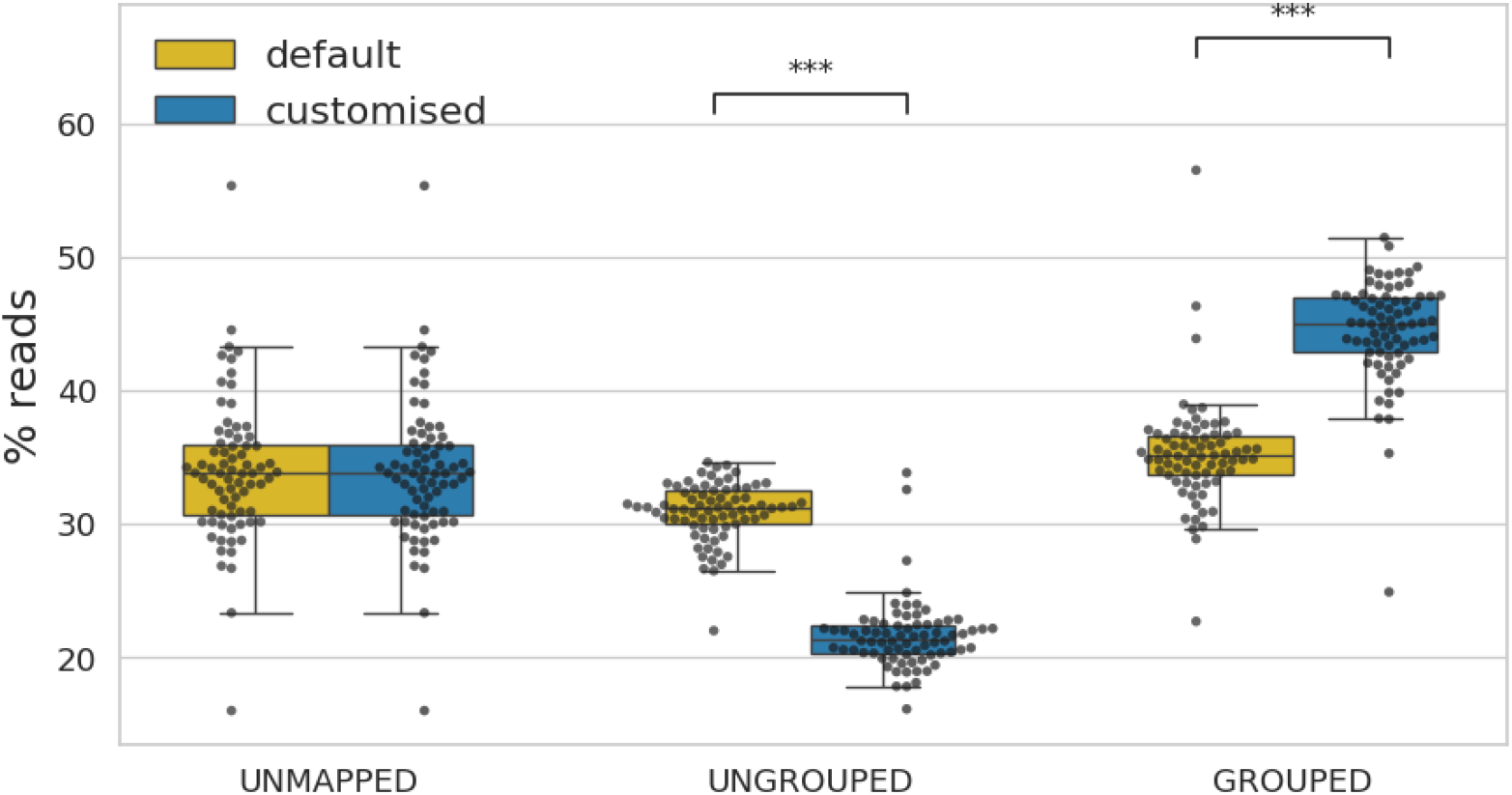
Impact of GPR mapping approach on functional profile. Distribution of metagenomic reads across functional categories comparing HUMAnN3 outcomes: default vs our comprehensive mapping approach. Our approach increases reads assigned to metabolic functions from approximately 35% to 45%, reducing the ungrouped fraction without changing gene-level detection. UNMAPPED: reads not aligned to gene families; UNGROUPED: reads aligned to gene families but not assigned to metabolic reactions; GROUPED: reads aligned to gene families and assigned to metabolic reactions. Statistical significance assessed by paired Wilcoxon signed-rank test (p < 0.001).

In terms of absolute number of reactions, the default output identifies ≈ 4,000 reactions in the human gut metagenomic dataset and our GPR mapping approach identifies ≈ 400,000 reactions in the same dataset for the same gene families i.e., no changes were made at gene level rather the difference is due to mapping of genes to function alone. However, comparing the absolute number of reactions across the two approaches is not reasonable due to different databases used as described in section 3.3. Therefore, all the reaction level comparison in this study is based on percentage rather than absolute numbers.

In addition, reaction prevalence across samples increased substantially, as shown in Figure 4. Vhile only 35% of reactions were core, meaning they were identified in all samples in default HUMAnN3 output, with the improved GPR mapping more than 80% of reactions were core. The low prevalence was due to missing GPR mapping giving a false impression that there was difference in core reaction profile. This directly addresses the data sparsity challenge that limits statistical power and hinders translational applications [2, 3].

**Figure 4:**
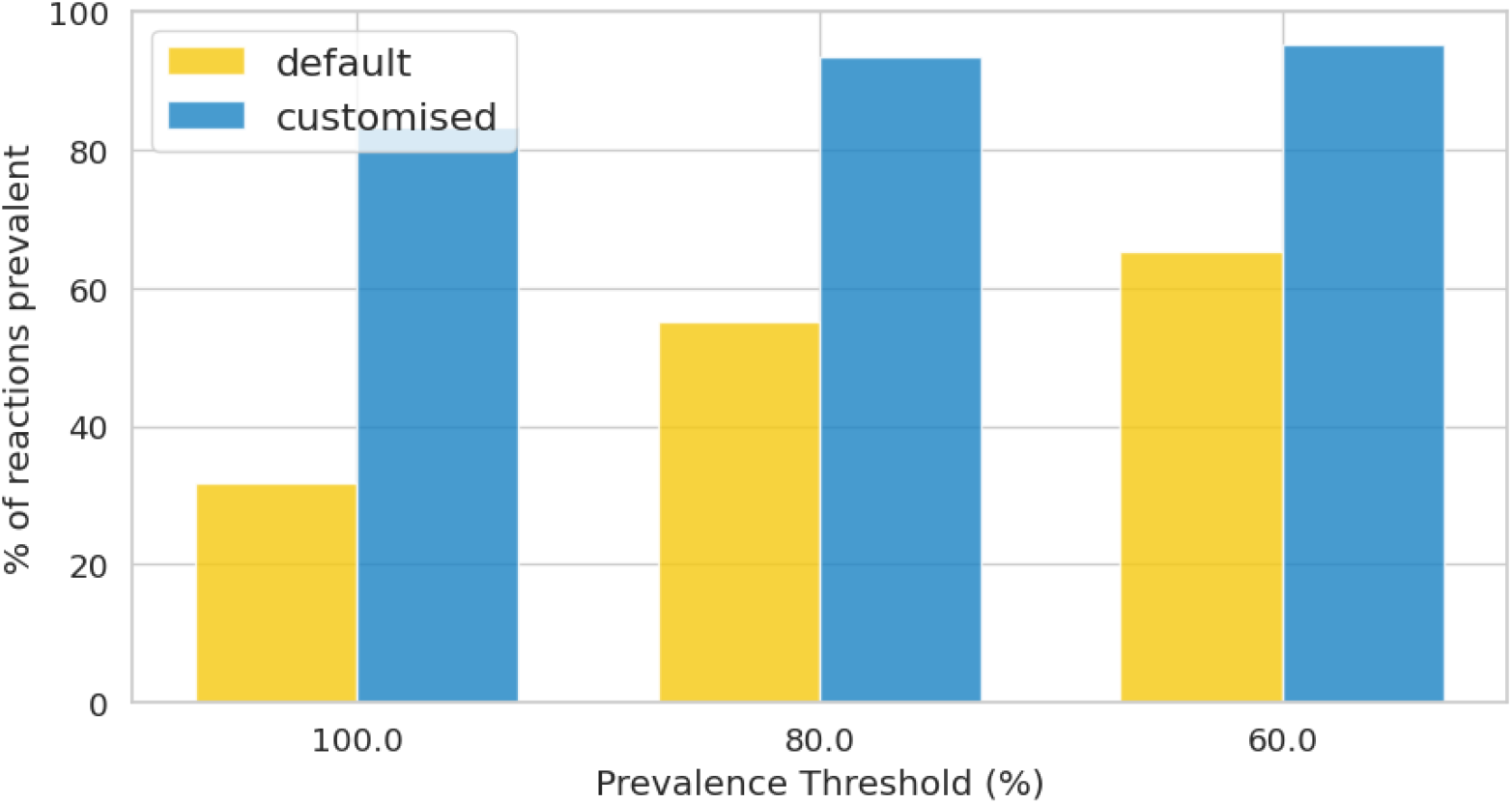
Comprehensive GPR mapping reduces data sparsity by increasing reaction prevalence across samples. Comparison of reaction prevalence between HUMAnN3 outcomes: default vs our comprehensive mapping approach. With default mapping, only 35% of reactions were core (detected in all samples), whereas our improved GPR mapping increased core reactions to over 80%.

### 3.3 Study Limitations

BioCyc comprises 21,244 strain-specific metabolic databases, including MetaCyc, and collectively covers metabolic reactions across all domains of life. MetaCyc serves as a well-curated reference database and therefore exhibits fewer inconsistencies than the broader BioCyc collection. However, MetaCyc is not free from inconsistencies, particularly in reaction identifiers, which can result in the same biochemical reaction being counted multiple times and thus inflate estimates of unique reaction numbers. Our mapping approach leverages the full BioCyc resource to achieve broader functional coverage than MetaCyc-based pipelines such as HUMAnN3, but consequently inherits a higher level of reaction-naming inconsistency and therefore, overestimation of number of reactions, relative to MetaCyc.

Additionally, Pfam-based mapping increases sensitivity but introduces potential false positives when domains are shared across distinct enzyme families. We mitigate this through hierarchical mapping (EC first) and provide modular files enabling users to balance sensitivity and specificity.

### 3.4 Future Directions and Broader Implications

Vhile our approach substantially increases reads mapped to metabolic functions, a considerable fraction of metagenomic reads still remain unmapped (reads not mapped to genes) or ungrouped (genes not mapped to metabolic function) (Figure 3). Presence of these groups is biologically plausible and complete elimination is neither expected nor realistic, due to non-coding regions, sequencing artifacts, and non-metabolic genes. However, a further systematic investigation is needed to ensure functions are not overlooked due to annotation gaps. While functional metagenomics pipelines remain constrained by cross-database integration, periodic remapping will be essential as curation improves, particularly with AI-based annotation advances.

Reaction-resolution profiling is more insightful than pathway-level resolution, but it generates high-dimensional datasets that challenge existing analytical paradigms designed for pathway-level or low-dimensional datasets. Specialised frameworks are needed in the field to handle data dimensionality, tailored to metabolic network topology. As microbiome datasets continue to grow in scale and complexity, such methodological innovations will be essential for extracting biological insights and enhancing translation potential.

Most critically, microbiome research requires fostering methodological introspection. Researchers must examine not only *what* tools output but *how* outputs are generated, including database dependencies, thresholds, and embedded assumptions. Microbiome science is inherently interdisciplinary; real translational impact requires collaborative teams bridging biological and computational domains to jointly diagnose limitations and co-develop solutions.

## 4 Conclusion

Comprehensive functional characterisation is essential for translating microbiome research into clinical and biotechnological applications, yet current approaches systematically underestimate metabolic capabilities. This work demonstrates that systematic integration of comprehensive reaction databases (BioCyc) with gene families (UniRef90) through complementary EC and Pfam strategies significantly increases functional coverage and prevalence across samples. These gains are derived from database integration alone, without predictive algorithms, demonstrating that substantial functional dark matter and data sparsity in microbiome studies are due to methodological artifacts.

By addressing one tractable yet systematically overlooked limitation, cross-database integration, this work represents a high-impact approach to advancing microbiome research and its translational potential. However, technical solutions alone are insufficient. Realising translational potential requires cultural shifts toward methodological introspection, moving beyond over-reliance on pathways as metrics of microbiome metabolic function to embrace multi-resolution profiling where researchers interrogate functional potential at resolutions appropriate to their biological questions and also critically examine the technical foundations that underpin the biological interpretation.

## 5 Data and Code Availability

All mapping files and analysis scripts are available from the FuncDarkMatt repository at https://github.com/quadram-institute-bioscience/FuncDarkMatt/ upon request. The repository includes Python scripts for database parsing and mapping, along with documentation for reproducing the workflow.

## 6 Funding

This work was supported by the Quadram Institute Small Pilot Project Grant. D.S. is funded by the QIB Early Career Fellowship Scheme. S.T. and A.T. are funded by BBSRC Core Capability Grant BB/CCG1860/1.

## 7 Author Contributions

Conceptualization, DS; methodology, DS and SKT; investigation, DS and SKT; formal analysis, DS and SKT; writing-original draft preparation, DS; writing-review and editing, DS, SKT and AT; supervision, DS; funding acquisition, DS and AT.

## 8 Acknowledgments

We thank John Wain for initiating discussions about limitations in microbiome research, and Alise Ponsero for valuable technical discussions.

## 9 Author Declarations

All authors have read and approved the manuscript. This manuscript has not been submitted, accepted or published elsewhere.

## 10 Competing Interests

The authors declare no competing interests.

## Notes

### Competing Interest Statement

The authors have declared no competing interest.

## References

[1] Alexandre Almeida, Stephen Nayfach, Miguel Boland, Francesco Strozzi, Martin Beracocliea, Zhou Jason Shi, Katherine S Pollard, Ekaterina Sakharova, Donovan H Parks, Philip Hugenholtz, Nicola Segata, Nikos C Kyrpides, and Robert D Finn. A unified catalog of 204,938 reference genomes from the human gut microbiome. Nature Biotechnology, 39(1): 105–114, 2021.

[2] Jason Lloyd-Price, Cesar Arze, Ashwin N Ananthakrishnan, Melanie Schirmer, Julian Avila-Pacheco, Tiffany W Poon, Elizabeth Andrews, Nadim J Ajami, Kevin S Bonham, Colin J Brislawn, David Casero, Holly Courtney, Antonio Gonzalez, Thomas G Graeber, A Brantley Hall, Kathleen Lake, Carol J Landers, Himel Mallick, Damian R Plichta, Mahadev Prasad, Gholamali Rahnavard, Jenny Sauk, Dmitry Shungin, Yoshiki Vázquez-Baeza, Richard A White, IBDMDB Investigators, Jonathan Braun, Lee A Denson, Janet K Jansson, Rob Knight, Subra Kugathasan, Dermot P B McGovern, Joseph F Petrosino, Thaddeus S Stappenbeck, Harland S Winter, Clary B Clish, Eric A Franzosa, Hera Vlamakis, Ramnik J Xavier, and Curtis Huttenhower. Multi-omics of the gut microbial ecosystem in inflammatory bowel diseases. Nature, 569(7758):655–662, 2019.

[3] Anna Heintz-Buschart and Paul Wilmes. Human gut microbiome: function matters. Trends in Microbiology, 26(7):563–574, 2018.

[4] David Zeevi, Tal Korem, Niv Zmora, David Israeli, Daphna Rothschild, Adina Weinberger, Orly Ben-Yacov, Dar Lador, Tali Avnit-Sagi, Maya Lotan-Pompan, Jotham Suez, Jemal Ali Mahdi, Elad Matot, Gal Malka, Noa Kosower, Michal Rein, Gili Zilberman-Schapira, Lenka Dohnalovä, Meirav Pevsner-Fischer, Rony Bikovsky, Zamir Halpern, Eran Elinav, and Eran Segal. Personalized nutrition by prediction of glycemic responses. Cell, 163(5):1079–1094, 2015.

[5] The Human Microbiome Project Consortium. Structure, function and diversity of the healthy human microbiome. Nature, 486(7402) :207–214, 2012.

[6] Eric A Franzosa, Lauren J McIver, Gholamali Rahnavard, Luke R Thompson, Melanie Schirmer, George Weingart, Karen Schwarzberg Lipson, Rob Knight, J Gregory Caporaso, Nicola Segata, and Curtis Huttenhower. Species-level functional profiling of metagenomes and metatranscriptomes. Nature Methods, 15(11):962–968, 2018.

[7] Sahar Abubucker, Nicola Segata, Johannes Goll, Alyxandria M Schubert, Jacques Izard, Brandi L Cantarel, Beltran Rodriguez-Mueller, Jeremy Zucker, Mathangi Thiagarajan, Bernard Henrissat, Owen White, Scott T Kelley, Barbara Methé, Patrick D Schloss, Dirk Gevers, Makedonka Mitreva, and Curtis Huttenhower. Metabolic reconstruction for metagenomic data and its application to the human microbiome. PLoS Computational Biology, 8(6):e1002358, 2012.

[8] Ines Thiele, Swagatika Sahoo, Almut Heinken, Johannes Hertel, Laurent Heirendt, Maike K Aurich, and Ronan MT Fleming. Personalized whole-body models integrate metabolism, physiology, and the gut microbiome. Molecular Systems Biology, 16(5):e8982, 2020.

[9] Sofia K Forslund, Rima Chakaroun, Maria Zimmermann-Kogadeeva, Lajos Markó, Judith Aron-Wisnewsky, Trine Nielsen, Lucas Moitinho-Silva, Trine S B Schmidt, Gwen Falony, Sara Vieira-Silva, Solia Adriouch, Renato Alves, Karen Assmann, Jean-Philippe Bastard, Till Birkner, Robert Caesar, Julien Chilloux, Luis Pedro Coelho, Leopold Fezeu, Nathalie Galleron, Gerard Helft, Richard Isnard, Boyang Ji, Michael Kuhn, Emmanuelle Le Chatelier, Antonis Myridakis, Lisa Olsson, Nicolas Pons, Edi Prifti, Benoit Quinquis, Hugo Roume, Joe-Elie Salem, Nataliya Sokolovska, Valentina Tremaroli, Mireia Valles-Colomer, Christian Lewinter, Nadja Sqndertoft, Helle Krogh Pedersen, Tue Hansen, Jens Peter Gqtze, Lars Kqber, Henrik Vestergaard, Torben Hansen, Jean-Daniel Zucker, Serge Hercberg, Jean-Michel Oppert, Ivica Letunic, Jens Nielsen, Fredrik Bäcklied, S Dusko Ehrlich, Marc-Emmanuel Dumas, Jeroen Raes, Oluf Pedersen, Karine Clément, Michael Stumvoll, Peer Bork, and MetaCardis Consortium. Combinatorial, additive and dose-dependent drug-microbiome associations. Nature, 600(7889):500–505, 2021.

[10] Jiwoong Kim, Min Soo Kim, Andrew Y. Koh, Yang Xie, and Xiaowei Zhan. Fmap: Functional mapping and analysis pipeline for metagenomics and metatranscriptomics studies. BMC Bioinformatics, 17:420, 10 2016.

[11] Peter D Karp, Richard Billington, Ron Caspi, Carol A Fulcher, Mario Latendresse, Anamika Kothari, Ingrid M Keseler, Markus Krummenacker, Peter E Midford, Quang Ong, Wai Kit Ong, Suzanne M Paley, and Pallavi Subhraveti. The biocyc collection of microbial genomes and metabolic pathways. Briefings in Bioinformatics, 20(4):1085—1093, 2019.

[12] Baris E Suzek, Yuqi Wang, Hongzhan Huang, Peter B McGarvey, Cathy H Wu, and UniProt Consortium. Uniref clusters: a comprehensive and scalable alternative for improving sequence similarity searches. Bioinformatics, 31 (6) :926–932, 2015.

[13] Jaina Mistry, Sara Chuguransky, Lowri Williams, Matloob Qureshi, Gustavo A Salazar, Erik LL Sonnhammer, Silvio CE Tosatto, Lisanna Paladin, Shriya Raj, Lorna J Richardson, Robert D Finn, and Alex Bateman. Pfam: The protein families database in 2021. Nucleic Acids Research, 49(D1):D412–D419, 2021.

[14] UniProt Consortium. Uniprot: the universal protein knowledgebase in 2021. Nucleic Acids Research, 49(D1):D480–D489, 2021.

[15] Francesco Beghini, Lauren J McIver, Aitor Blanco-Míguez, Leonard Dubois, Francesco Asnicar, Sagun Maharjan, Ana Mailyan, Paolo Manghi, Matthias Scholz, Andrew Maltez Thomas, Mireia Valles-Colomer, George Weingart, Yancong Zhang, Moreno Zolfo, Curtis Huttenhower, Eric A Franzosa, and Nicola Segata. Integrating taxonomic, functional, and strain-level profiling of diverse microbial communities with biobakery 3. eLife, 10:e65088, may 2021.

